# Quality assurance of hematopoietic stem cells by macrophages determines stem cell clonality

**DOI:** 10.1101/2022.03.07.483327

**Authors:** Samuel J. Wattrus, Mackenzie L. Smith, Cecilia Pessoa Rodrigues, Elliott J. Hagedorn, Bogdan Budnik, Leonard I. Zon

**Author notes:** Section of Hematology and Medical Oncology and Center for Regenerative Medicine, Boston University School of Medicine and Boston Medical Center, Boston, MA, USA.

## Abstract

Tissue-specific stem cells persist for a lifetime and can differentiate to maintain homeostasis or transform to initiate cancer. Despite their importance, there are no described quality assurance mechanisms for newly formed stem cells. We observed intimate and specific interactions between macrophages and nascent blood stem cells in zebrafish embryos. Macrophage interactions led to two outcomes — removal of cytoplasmic material and stem cell division, or complete engulfment and stem cell death. Stressed stem cells were marked by surface Calreticulin, which stimulated macrophage interactions. Using cellular barcoding, we found that calreticulin knock-down or embryonic macrophage depletion reduced the number of stem cell clones that established adult hematopoiesis. Our work supports a model in which embryonic macrophages determine hematopoietic clonality by monitoring stem cell quality.

## Main Text

Tissue stem cells born during embryogenesis support homeostasis for life. Despite the importance of these cells for proper tissue function, there are no described mechanisms of quality assurance for newly formed stem cells. To explore this possibility, we studied zebrafish embryonic blood development. Hematopoietic stem and progenitor cells (HSPCs) emerge from the ventral wall of the dorsal aorta (VDA), enter circulation, and lodge in the embryonic niche - a vascular plexus called the caudal hematopoietic tissue (CHT) [1, 2]. HSPCs rapidly expand in the CHT for a few days before migrating to the kidney marrow, the adult hematopoietic niche. In the niche, HSPCs interact with a variety of cell types, including vascular endothelial cells, mesenchymal stromal cells, and macrophages [3–6]. *In vivo* clonal labeling studies show that 20 – 30 of the hematopoietic stem cell (HSC) clones born in the VDA ultimately give rise to the adult blood system [7]. It remains unclear if nascent HSCs from the VDA undergo quality assurance before establishing adult hematopoiesis. Here, using live imaging and cellular barcoding, we found discrete interactions between stem cells and embryonic macrophages that regulated the number of long-lived hematopoietic stem cells clones that produce blood in adulthood.

## Results

### Macrophages interact with nascent HSPCs in the niche

Macrophages help maintain homeostasis by modulating inflammation, producing cytokines, and patrolling to clear dead, stressed, or aged cells [8–10]. Given these roles in somatic tissue, the enrichment of macrophages in the CHT, and previous observations between macrophages and hematopoietic cells [6], we investigated macrophage function in the niche. We undertook high-resolution live imaging using *mpeg1:mCherry;runx1+23:EGFP* zebrafish embryos with mCherry^+^ macrophages and EGFP^+^ HSPCs [4, 11]. Shortly after lodgment in the CHT, HSPCs were physically contacted by a nearby macrophage, their surfaces were scanned, and fluorescent material was taken up by the macrophage (Fig. 1A; Movie S1). From 56 to 106 hours post-fertilization (hpf), approximately 20-30% of HSPCs in the niche were engaged by macrophages at any one time (Fig. 1B). These interactions were specific to HSPCs; macrophage engagement with erythrocytes and endothelial cells was significantly lower (0.6 - 3.9% of erythrocytes and 0.5 – 6.7% of endothelial cells) (Fig. S1A). Macrophages physically contacted HSPCs for up to 45 minutes and sometimes extracted fluorescent HSPC material. We classified interactions with fluorescence transfer into two types: “dooming” during which the HSPC was fully engulfed and destroyed by the macrophage, or “grooming” during which the HSPC was left intact and had a small portion of cellular material taken up by the macrophage (Movie S2). Similar interactions were identified in other zebrafish HSPC-reporter lines *cd41:GFP* and *cmyb:GFP* (Figs. S1D-E). To examine if macrophage-HSPC interactions occurred in mammals, we studied E14.5 murine fetal liver sections by immunofluorescence and found that on average 33% of c-Kit^+^ hematopoietic cells were in contact with F4/80^+^ macrophages. These contacts included c-Kit^+^ cells being pinched or fully engulfed by macrophages, similar to our observations in zebrafish (Fig. S1F). Overall, these data identify novel macrophage-HSPC interactions in the embryonic hematopoietic niche.

**Fig. 1.**
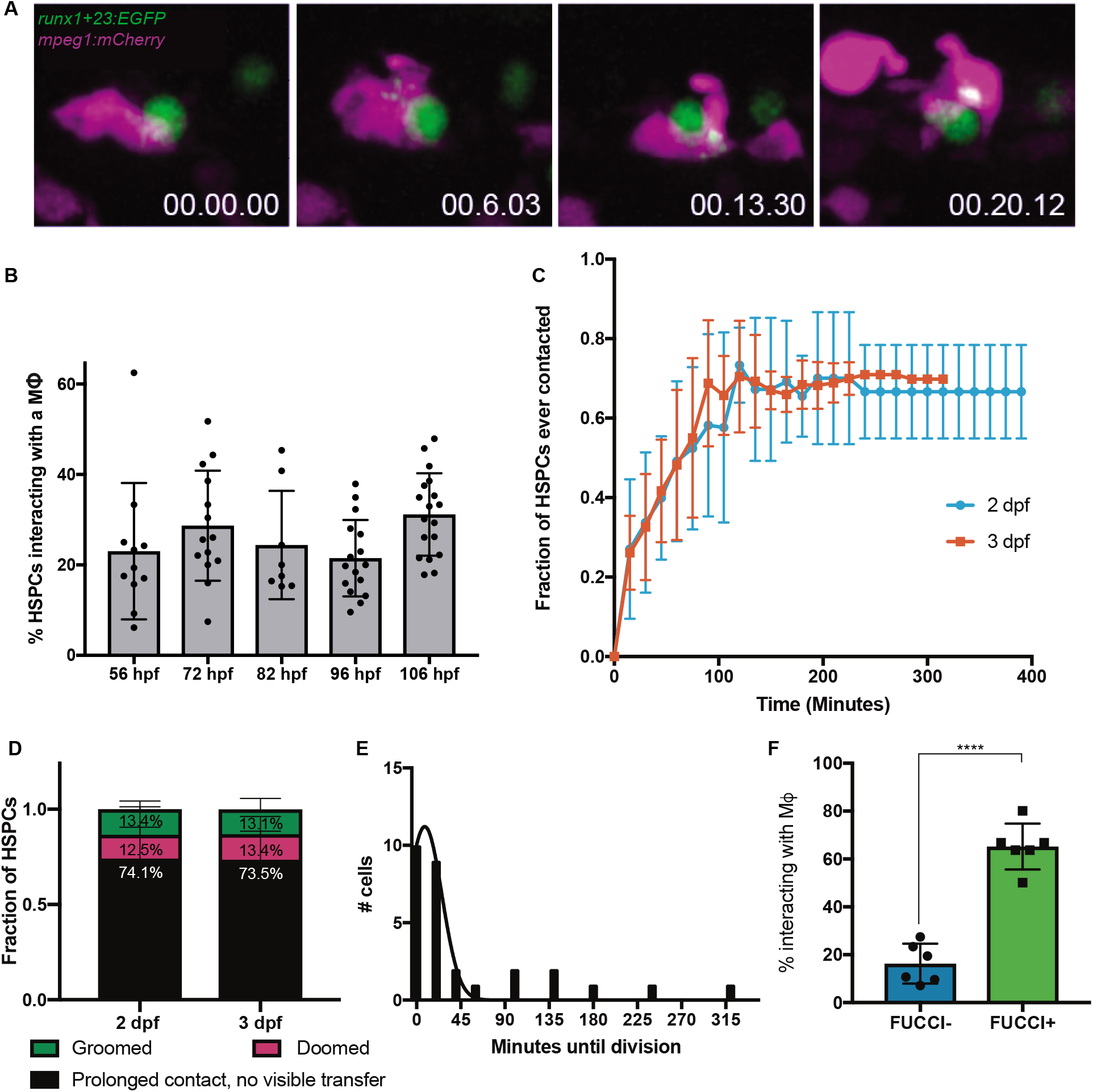
Macrophages make intimate interactions with newly formed HSPCs. (**A**) Time lapse live-imaging identifies prolonged cell-cell contacts between *runx1+23:EGFP*^+^ HSPCs and *mpeg1:mCherry*^+^ primitive macrophages in which cytoplasmic fluorescent material is exchanged. Time is given in hh:mm:ss. (**B**) Approximately 20-30% of HSPCs interact with macrophages in the CHT at any one time from 56 hpf to 106 hpf. Mean +/− s.d. (**C**) High-resolution cell tracking of individual *runx1+23:mCherry^+^* cells over several hours in the CHT reveals that the majority of HSPCs eventually make sustained contact with macrophages (> 5 minutes). Mean +/− s.d. (**D**) Approximately 26% of contacted cells are either groomed or doomed. Mean +/− s.d. (**E**) HSPCs frequently complete a cell division shortly after macrophage interactions. Approximately 81% of HSPCs complete a division within 30 minutes of interacting with a macrophage. Mean +/− s.d. (**F**) Around 65% of Fucci^+^ HSPCs in G2/M phase interact with macrophages at any one time, as compared to less than 20% of Fucci^−^ HSPCs. Mean +/− s.d., Unpaired t test; ****P<0.0001.

To better characterize macrophage-HSPC interactions, we tracked individual HSPCs in the CHT over 4-6 hours and recorded all interactions with macrophages at 2 or 3 days post-fertilization (dpf). We found that 70% of HSPCs eventually experienced prolonged macrophage contact (Fig. 1C). Of these interactions, 13% of HSPCs were groomed, and 13% were fully engulfed and doomed (Fig. 1D). Interestingly, 81% of HSPC divisions occurred within 30 minutes following macrophage contact (Fig. 1E). Using the *Tg(EF1a:mAG-zGem(1/100))^rw0410h^* (Fucci) transgene [12] that labels cells in G2M phase of the cell cycle, we found that approximately 65% of Fucci^+^ HSPCs contacted macrophages, compared to only 16% of Fucci^−^ HSPCs (Fig. 1F). We next assessed the viability of HSPCs that were engulfed by macrophages. Staining for cell death *in vivo* with Acridine Orange or an Annexin V-YFP construct [13] showed almost no apoptotic HSPCs in the CHT at baseline (Figs. S1B-C). Only after full engulfment by a macrophage did HSPCs exhibit apoptosis (Movie S3). Taken together, these data identify a set of prolonged macrophage-HSPC interactions that either precede HSPC division or death.

### A subset of primitive macrophages regulates stem cell clone number

As we saw proliferation following macrophage-HSPC interactions, we next sought to determine if this might have an effect on the number of stem cell clones that contribute to adult hematopoiesis. We used TWISTR (tissue editing with inducible stem cell tagging via recombination) [14] to combine alteration of embryonic gene expression by morpholino injection with color labeling of HSCs by Zebrabow. *Zebrabow-M;draculin:CreER^T2^* embryos allow for unique lineage labeling of individual HSC clones at 24 hpf (Fig. 2A) [7, 15]. To deplete embryonic macrophages, we injected the *irf8* morpholino into *Zebrabow-M;draculin:CreER^T2^* embryos to block macrophage differentiation from the one-cell stage [16] or delivered clodronate liposomes to deplete macrophages at various timepoints: 28 hpf, just prior to HSPC emergence in the VDA, 48 hpf, before HSPC lodgment in the CHT, and 72 hpf, after HSPCs have lodged in the CHT. Analysis of adult marrow myelomonocytes from macrophage-depleted animals revealed a consistent reduction in hematopoietic clonality compared to sibling controls for all timepoints (Fig. 2B). These results demonstrate that embryonic macrophages regulate long-term HSC clone number after VDA emergence.

**Fig. 2.**
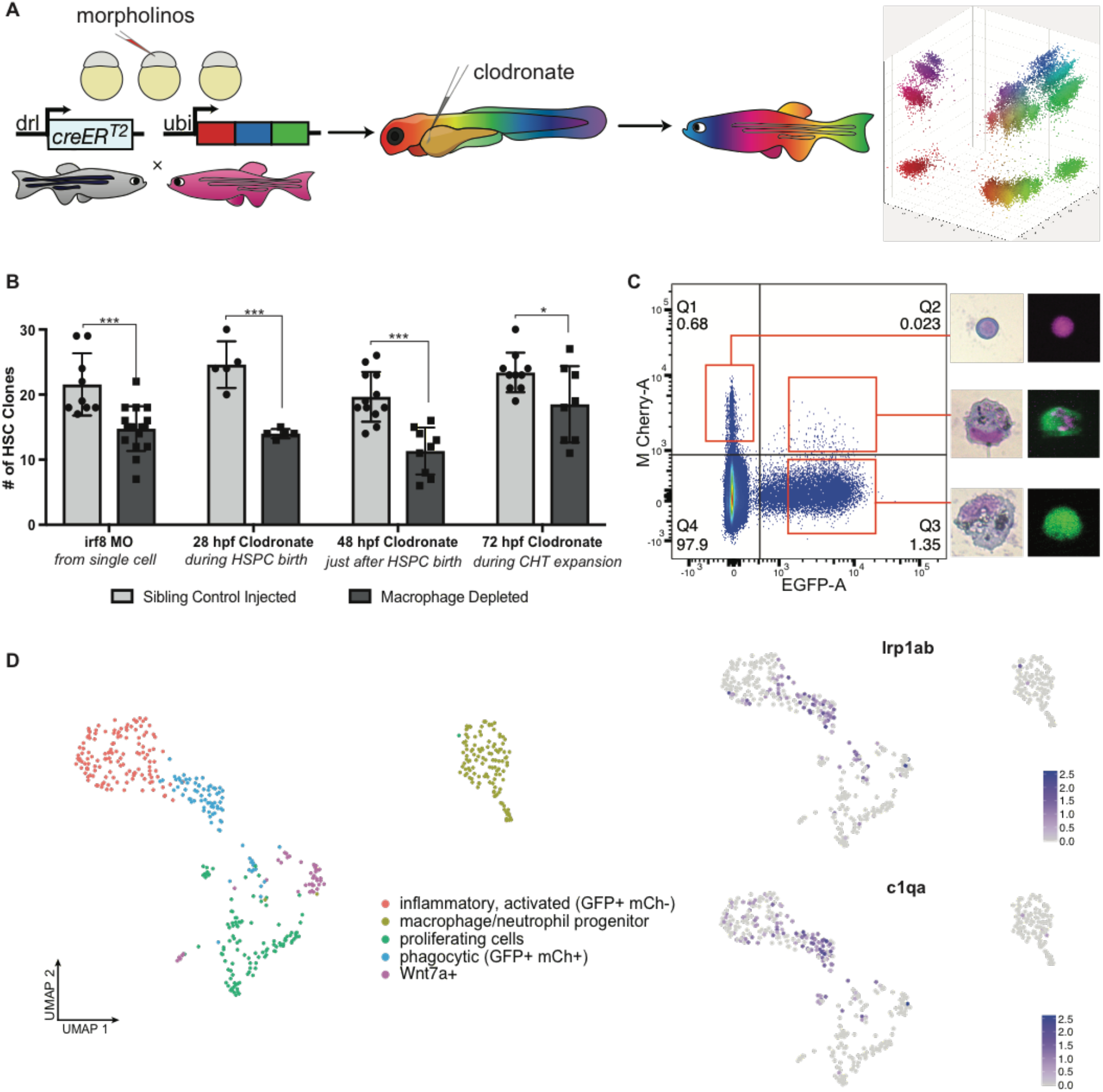
Macrophages in the CHT regulate stem cell clonality. (**A**) A schematic overview of the *Zebrabow-M* system: animals with 15-20 insertions of tandem multicolor fluorescent cassettes are crossed to a *draculin:CreER^T2^* line, which enables specific clonal labeling of the lateral plate mesoderm lineage. By treating animals with 4-OHT at 24 hpf just after HSC specification, individual stem cells express unique fluorescent hues. These unique hues can be identified in adult marrow and quantified to determine the number and size of stem cell clones. (**B**) Families of *Zebrabow-M;draculin:CreER^T2^* animals injected with either clodronate liposomes or the *irf8* morpholino exhibit reduced numbers of HSC clones in the adult marrow, even when macrophages are not depleted until after emergence from the VDA. Mean +/− s.d., Unpaired t test; *P<0.05, ***P<0.001. (**C**) Macrophages (*mpeg1:EGFP*^+^) which have interacted with HSPCs (*runx1+23:mCherry*^+^) and removed fluorescent material can be identified and harvested by flow cytometry. (**D**) Uniform Manifold Approximation and Projection (UMAP) plots shows that macrophages which engage HSPCs are marked by *lrp1ab* and *c1qa*. Spectral scale reports z-scores.

To better understand the mechanism and cellular consequences of macrophage-HSPC interactions observed in the CHT, we pursued transcriptomic analysis of niche macrophages. Because macrophages can take up fluorescent material from HSPCs, we reasoned it would be possible to identify interacting macrophages by their fluorescence profile. Indeed, flow cytometry of dissociated *mpeg1:EGFP;runx1+23:mCherry* embryos revealed a rare population of EGFP^+^mCherry^+^ cells (1.67% of all EGFP^+^ cells), which were morphologically consistent with macrophages containing small HSPC fragments (Fig. 2C). We dissected embryonic zebrafish tails at 72 hpf and purified interacting macrophages (EGFP^+^mCherry^+^) and non-interacting macrophages (EGFP^+^mCherry^−^) for plate-based single-cell mRNA sequencing. By recording single-cell indices and combining FACS parameters with single-cell transcriptomic profiles, we identified a single population of macrophages which segregated by both gene expression profile and mCherry fluorescence (Fig. 2D). These cells exhibited a relative enrichment for genes associated with engulfment, lysosomal degradation, and cholesterol transporter activity and were marked by genes including *hmox1a, ctsl.1, slc40a1, lrp1ab,* and *c1qa* (Fig. 2D; Fig. S2A) [17]. We validated these data through injection of a fluorescent cholesterol mimic, treatment with Lysotracker dyes, and *in situ* hybridization (Figs. S2B-D). Together, these data show that a transcriptionally distinct and relatively homogenous subset of macrophages engage HSPCs in the CHT for both grooming and dooming behaviors.

### Surface Calreticulin drives macrophage-HSPC interactions

To gain insight into the proteinaceous material taken up by macrophages, we pursued few-cell proteomics [18] to compare interacting to non-interacting macrophages isolated from 72 hpf embryos. We identified 203 peptides enriched in interacting macrophages potentially representing a repertoire of macrophage proteins involved in the process of macrophage-HSPC interaction or HSPC-derived proteins taken up during the interaction. To identify molecular patterns recognized on HSPCs, we excluded peptides with enriched transcripts in interacting macrophages and compared the remaining peptides to the Cell Surface Protein Atlas [19]. Notably, cell surface peptides enriched in EGFP^+^mCherry^+^ macrophages included three Calreticulin paralogs: *calr, calr3a,* and *calr3b* (Fig. 3A). Though Calreticulin typically functions as a calcium-binding chaperone protein in the endoplasmic reticulum, it can also be translocated and displayed on the cell surface as an “eat me” signal [9, 10, 20]. Based on our proteomic results, we hypothesized that HSPCs display surface Calreticulin leading to macrophage interaction, and that increasing levels of Calreticulin could lead to grooming and dooming. We found that 30% of *runx1+23^+^* HSPCs at 72 hpf exhibited classic punctate surface Calr staining [21] (Fig. 3B), similar to the percentage of HSPCs we found interacting with macrophages at this time in the CHT (Fig. 1B). Additionally, the canonical surface Calreticulin binding partners, *lrp1ab* and *c1qa,* were enriched in interacting macrophages by single-cell mRNA-seq (Fig. 2D). Together with complement C1q subcomponent A (*c1qa*), Lrp1ab contacts Calreticulin and forms a bridging complex to initiate phagocytic activity [10, 21, 22]. These results show that Calreticulin decorates the surface of a subset of HSPCs and may signal for macrophage interaction.

**Fig. 3.**
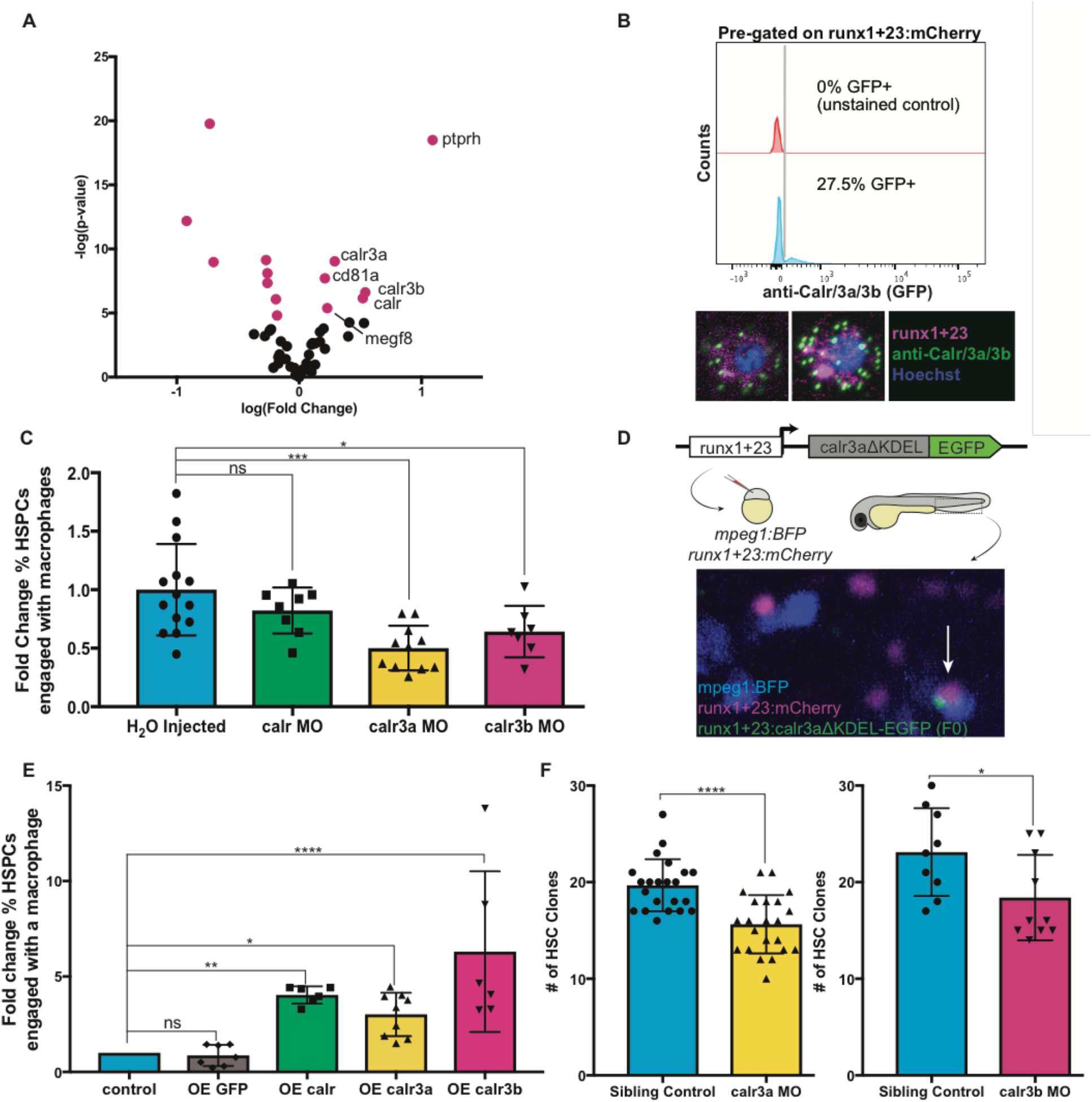
Calreticulin drives HSPC-macrophage interactions to regulate clonality. (**A**) Hundreds of interacting and noninteracting macrophages were sorted and processed for few-cell proteomics by LC-MS/MS. Analysis of differentially enriched putative surface proteins from interacting macrophages identifies three paralogs of Calreticulin. Magenta dots indicate differentially enriched surface proteins. (**B**) Antibody staining and flow cytometry identifies ~30% of live *runx1+23:mCherry*^+^ HSPCs stain positive for Calreticulin. Confocal imaging displays punctate surface stain, similar to previous reports. (**C**) Knock-down of *calr3a* or *calr3b* by morpholino injection significantly reduces the fraction of HSPCs interacting with macrophages at any one time. Data are normalized to H_2_O injected embryos. Mean +/− s.d., One-way ANOVA with Dunnett’s multiple comparisons test; *P<0.05, ***P<0.001. (**D**) Schematic and example of overexpression approach. Calreticulin paralogs lacking the ER-retention KDEL sequence were fused to EGFP, driven by the HSPC-specific *runx1+23* enhancer, and injected into stable *runx1+23:mCherry;mpeg1:BFP* embryos to generate mosaic animals to compare HSPCs overexpressing Calreticulin against unperturbed HSPCs in the same embryo. Arrow indicates an HSPC overexpressing *calr3a* engaged by a macrophage. (**E**) HSPCs overexpressing *calr, calr3a*, or *calr3b* are more frequently engaged by macrophages compared to non-overexpressing HSPCs in the same embryos. Overexpressing GFP alone has no effect on macrophage engagement. Data are normalized to the fraction of non-overexpressing HSPCs in each individual animal. Mean +/− s.d., One-way ANOVA with Dunnett’s multiple comparisons test; *P<0.05, **P<0.01, ****P<0.0001. (**F**) Inhibition of *calr3a* or *calr3b* by morpholino injection reduces the number of HSC clones that contribute to adult marrow compared to control injected embryos from the same family. Mean +/− s.d., Unpaired t test; *P<0.05, ****P<0.0001.

To study the role of Calreticulin in macrophage-HSPC interactions, we used morpholinos to knock-down gene expression and quantified these interactions. We observed a significant reduction in the percentage of HSPCs engaged by macrophages after either *calr3a* or *calr3b* knock-down (Fig. 3C). This effect was reversed in genetic rescue experiments (Figs. S3A-B). We generated parabiotic fusions to join circulation of embryos with or without calreticulin knock-down and found that HSPCs with *calr3a* knock-down had reduced interactions with unperturbed macrophages, indicating that Calreticulin presentation is likely HSPC autonomous (Fig. S3C). We then tested the effect of a constitutively surface-translocated form of Calreticulin expressed under the HSPC specific *runx1+23* enhancer [4] (Fig. 3D). The mosaic nature of the expression of this construct allows simultaneous comparison of HSPCs with or without Calreticulin overexpression in the same embryo. HSPCs overexpressing either *calr, calr3a,* or *calr3b* were 3 to 5-fold more likely to interact with macrophages compared to unperturbed HSPCs in the same embryos (Fig. 3E). When *calr3a* or *calr3b* were knocked down, we found that the frequency of both grooming and dooming type interactions dropped, with a more severe decrease in dooming (Fig. S3D). Conversely, when levels of Calreticulin were elevated with overexpression constructs, nearly all expressing cells were doomed (Fig. S3E). Taken together, these data show that surface display of Calreticulin promotes macrophage-HSPC interactions in the CHT and suggest that differing levels of Calreticulin determine if an HSPC is groomed or doomed.

To determine if Calreticulin-dependent interactions during development were responsible for regulating HSC clone number into adulthood, we knocked down *calr3a* or *calr3b* during development and color labeled HSCs at 24 hpf. At 2 months, adult marrow analysis in these animals showed a significant reduction in the number of HSC clones compared to sibling controls (Fig. 3F). These data show that Calreticulin-dependent interactions in development support a greater number of long-lived HSC clones.

### Macrophages buffer HSPC stress and promote cell cycling

We next wanted to assess calreticulin function in HSPC development. To analyze changes to HSPC budding in the VDA we injected either the *irf8, calr3a,* or *calr3b* morpholino into *cd41:GFP;kdrl:mCherry* embryos, which permit visualization of the endothelial-to-hematopoietic transition [23]. Quantification of GFP^+^mCherry^+^ cells in the aortic floor from 32 – 46 hpf revealed no significant difference in HSPC budding with loss of macrophages or calreticulin (Fig. S4A). Serial imaging and quantification of *cd41:GFP^+^* cells in the CHT over early development revealed that knock-down of *calr3a* or *calr3b* did not affect HSPC numbers through 60 hpf, but later reduced HSPCs at 72 and 84 hpf (Figs. S4B-C). The loss of HSPCs in *irf8, calr3a,* and *calr3b* morphants was not due to apoptosis (Fig. S4D). Rather, depletion of macrophages or knock-down of *calr3a* or *calr3b* significantly reduced the fraction of proliferative HSPCs in the CHT at 72 hpf, as measured by EdU incorporation (Fig. 4A). This is in agreement with the association of macrophage-HSPC interactions with cell division identified by live imaging of the CHT (Figs. 1E–F). These data show that Calreticulin-dependent macrophage-HSPC interactions serve to expand and maintain HSPCs during early development by promoting proliferation in the CHT.

**Fig. 4.**
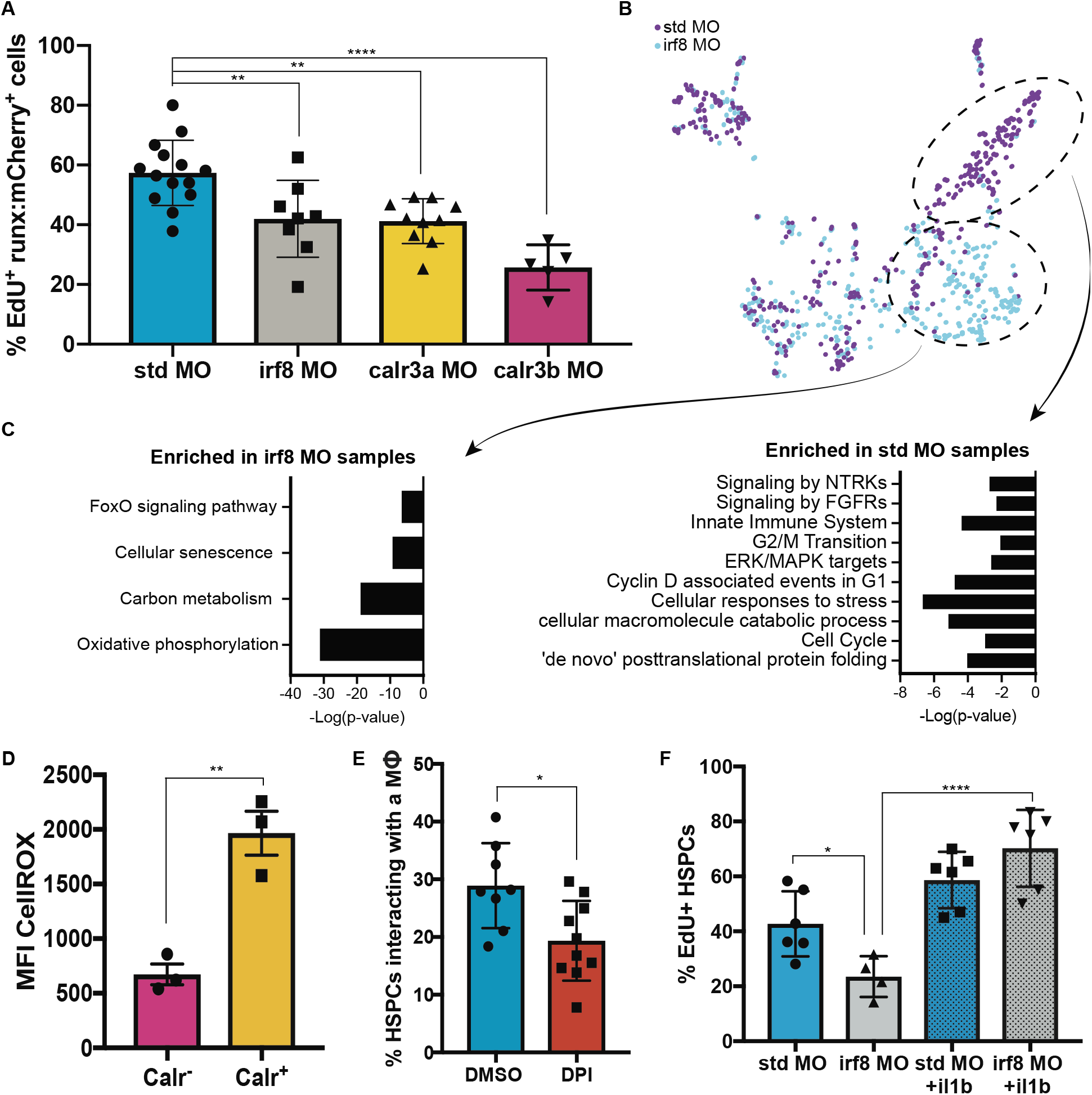
Macrophages buffer HSPC stress and regulate HSPC expansion. (**A**) EdU staining of *runx1+23:mCherry* embryos injected with either the *calr3a*, *calr3b*, or *irf8* morpholinos identifies a significant reduction in the fraction of proliferating HSPCs in the CHT at 3 dpf. Mean +/− s.d., One-way ANOVA with Dunnett’s multiple comparisons test; **P<0.01, ****P<0.0001. (**B**)(**C**) Single-cell mRNA-seq analysis of *runx1+23*^+^ FACS purified cells from 3 dpf embryos injected with either the *irf8* or non-targeting control morpholino. Subsets of HSPCs are enriched in the *irf8* or control sample. KEGG enrichment pathways reveals a population of stressed HSPCs that persist in the absence of macrophages and a population of cycling cells enriched in the control sample. (**D**) Embryonic HSPCs marked by surface Calreticulin exhibit higher levels of ROS as marked by CellROX intensity. (**E**) Inhibition of ROS by drug treatment with the NADPH oxidase inhibitor, diphenylene iodonium (DPI), significantly reduces the fraction of HSPCs engaged by macrophages in the CHT. Mean +/− s.d., Unpaired t test; *P<0.05. (**F**) Expression of *il1b* by heat shock is able to rescue the effect of macrophage depletion on HSPC proliferation. Mean +/− s.d., One-way ANOVA with Sidak’s multiple comparisons test; *P<0.05, ****P<0.001.

To molecularly evaluate the effect of macrophage interactions on HSPCs and the qualities of HSPCs that lead to surface Calreticulin, we injected *runx1+23:mCherry* embryos with the *irf8* or control morpholino and performed single-cell mRNA-seq on sorted HSPCs at 72 hpf. This analysis identified a subset of HSPCs enriched in *irf8* morphants marked by genes associated with FoxO activity and cellular senescence (Figs. 4B–C). FoxO activity is induced in response to elevated reactive oxygen species (ROS) and mediates detoxification of ROS and repair of ROS-induced damage [24, 25]. The enrichment of HSPCs with FoxO activity in *irf8* morphants suggested that there may be ROS accumulation in stem cells during embryogenesis which is ordinarily resolved by macrophages. In murine HSCs, FoxO deletion results in dysregulation of apoptosis, cell cycling, and colony formation due to resulting ROS accumulation [26]. In agreement with this, flow cytometric analysis identified higher levels of ROS in *runx1+23^+^* HSPCs marked by surface Calreticulin, with significant correlation between levels of ROS and surface Calreticulin (Spearman’s Correlation; ****P< 2.2e-16) (Fig. 4D). Treatment with the ROS inhibitor diphenylene iodonium reduced macrophage-HSPC interactions at 3 dpf (Fig. 4E). These results show that in the absence of macrophages, a population of HSPCs with elevated ROS accumulates, and this is associated with surface Calreticulin and subsequent macrophage interactions.

Embryos injected with the *irf8* morpholino also lost a subset of HSPCs marked by genes associated with cell cycling and ERK/MAPK signaling (Figs. 4B–C). In accordance with this, depleting macrophages or inhibiting ERK/MAPK activity from 48-72 hpf reduced HSPC proliferation without perturbing the frequency of macrophage-HSPC interactions. ERK/MAPK inhibition in the context of macrophage depletion did not further reduce proliferation, indicating that macrophages likely stimulate division via this pathway (Fig. S5A). As inflammation is a critical determinant of HSPC proliferation in development [27, 28], we reasoned that pro-proliferative cytokines produced by the macrophages, such as *il1b* and *tnfb*, could be responsible for the cell cycling phenotype (Fig. S5B) [28, 29]. To demonstrate this, we rescued HSPC proliferation after macrophage depletion by heat shock overexpression of *il1b* [28] (Fig. 4F). These results show that HSPC cycling in the CHT is mediated partly through ERK/MAPK and that signals produced by CHT macrophages including *il1b* can induce HSPC proliferation.

## Discussion

Our data support a model in which macrophages of the embryonic niche vet the quality of newly formed HSPCs through prolonged physical contact leading to either expansion or engulfment. This process is mediated by the display of Calreticulin on the cell surface, which is associated with elevated ROS. It has previously been reported that metabolic shifts during HSPC generation in the VDA elevate ROS to mediate HIF1α stabilization [30]. Cells with high levels of ROS are also at elevated risk for DNA damage and dysfunction [24, 25]. Our work suggests that while ROS generation in the VDA promotes stem cell emergence, titration of ROS levels is ultimately required for normal hematopoiesis. Although we see no evidence for *vcam1* expression in embryonic macrophages, previous imaging studies have indicated a role for macrophages in HSPC homing [6]. Macrophages are involved in HSPC mobilization in the zebrafish aorta [31], and murine macrophage subpopulations facilitate HSPC engraftment [32]. In contrast, our studies find that macrophages in the CHT remove stem cell clones with high surface Calreticulin which have not downregulated ROS production. Healthy HSPCs with low-to-moderate levels of ROS and Calreticulin experience prolonged macrophage contact, avoid complete engulfment, and receive pro-proliferative signals to enable competition for marrow colonization.

Our work establishes that stem cells are quality assured during development, and this impacts the clones that contribute to blood formation in adulthood. Calreticulin functions as an “eat me” molecule that initiates macrophage-HSPC interaction and leads to programmed cell clearance or stem cell expansion. Orthologs of CD47 and SIRPα, “don’t eat me” signals, have not been identified in zebrafish, but other primitive signals could participate in macrophage behaviors. This quality assurance mechanism may also be utilized in adulthood in response to environmental stress, such as during marrow transplantation or in clonal stem cell disorders including myelodysplasia and leukemia. Macrophages may selectively expand or remove clones of tissue-specific stem cells in other systems similar to our findings. Other tissue stem cells rely on macrophages to assure adequate tissue regeneration [33], and this could occur through selective proliferation of certain clones. Manipulating this quality assurance mechanism may have significant therapeutic implications for disorders of stem cell biology and tissue regeneration.

## Materials and Methods

### Animal models

Wild-type zebrafish AB, *casper* or *casper*-EKK, and transgenic lines *cd41*:*GFP* [34], *runx1+23:EGFP* [4], *runx1+23*:*mCherry* [*runx1+23*:*NLS-mCherry*] [4], c*myb:GFP* [35]*, kdrl*:*mCherry* [*kdrl*:*Hsa.hras-mCherry*] [36], *Zebrabow-M* [15] *draculin:CreER^T2^*[37], *mpeg1:EGFP* [11], *mpeg1:mCherry* [11]*, mpeg1:BFP* [*mpeg1:TagBFP*]*, LCR:EGFP* [38], *EF1a:mAG-zGem(1/100))^rw0410h^* [12], and *hsp70l:il1b* [28] were used in this study. Alternative gene names are listed in parenthesis and full transgene names are listed in brackets. Wildtype C57BL/6J mice (Jackson Labs stock #000664) were also used in this study. All animals were housed at Boston Children’s Hospital and handled according to approved Institutional Animal Care and Use Committee (IACUC) of Boston Children’s Hospital protocols.

### Single-cell mRNA-seq preparation

For single-cell mRNA-sequencing of embryonic macrophages, several hundred 72 hpf *runx1+23:mCherry;mpeg1:EGFP* embryos were anesthetized with tricaine overdosed and transected with a scalpel to harvest tail tissue. For analysis of HSPCs from *irf8* morpholino injected animals, 72 hpf *runx1+23:mCherry* embryos were anesthetized with tricaine and chopped with razor blades. Tissue was dissociated using Liberase (Roche) and cells were harvested by FACS using a BD FACSAria II (BD Biosciences). We used 3nM DRAQ7 (Abcam) for Live/Dead staining. Viable single cells were FACS sorted into 384-well plates, called cell capture plates, that were ordered from Single Cell Discoveries, a single-cell sequencing service provider based in the Netherlands. Each well of a cell capture plate contains a small 50 nl droplet of barcoded primers and 10 μl of mineral oil (Sigma M8410). After sorting, plates were immediately spun and placed on dry ice. Plates were stored at −80° C.

Plates were shipped on dry ice to Single Cell Discoveries, where single-cell mRNA sequencing was performed according to an adapted version of the SORT-seq protocol [39] with primers described in [40]. Cells were heat-lysed at 65° C followed by cDNA synthesis After second-strand cDNA synthesis, all the barcoded material from one plate was pooled into one library and amplified using in vitro transcription. Following amplification, library preparation was done following the CEL-Seq2 protocol [41] to prepare a cDNA library for sequencing using TruSeq small RNA primers (Illumina). The DNA library was paired-end sequenced on an Illumina Nextseq™ 500, high output, with a 1×75 bp Illumina kit (read 1: 26 cycles, index read: 6 cycles, read 2: 60 cycles)

### Single-cell mRNA-seq analysis

For each dataset, paired-end reads were aligned to the zebrafish transcriptome using BWA [42]. Data was demultiplexed as described in [43]. Mapping and generation of count tables were automated using the MapAndGo script. We corrected read counts for UMI barcodes by removing duplicate reads that had identical library, cellular, and molecular barcodes and mapped to the same gene. Transcript counts were adjusted using Poissonian counting statistics to yield the number of UMIs detected per cell. Counts were imported into R using the Seurat suite version 3.0 [44]. Low quality cells were filtered out by keeping only cellular barcodes with greater than 500 UMIs per cell, greater than 500 genes per cell, and less than 9 percent of reads mapping to mitochondrial genes. Data were log normalized and scaled, and the 2000 most variable genes were identified. We reduced dimensionality using PCA, selecting 15 PCs that explained the majority of variation in the data. PC loadings were used as input for a graph-based approach to cluster cells by type and as input for Uniform Manifold Approximation and Projection (UMAP). Differentially expressed genes were identified using a Wilcox Rank Sum test to compare count data between clusters. Significant genes were obtained using an FDR threshold of 0.05. To combine single cell transcriptional profiles with FACS profiles, individually sorted cell indices were merged with single cell RNA data by matching well ID.

For further analysis of purified *runx1+23:mCherry^+^* cells, we discarded the reads mapping to ERCC spike-ins and cells having transcripts correlating to mitochondrial and ribosomal genes with a Pearson’s correlation coefficient > 0.65. Next, RaceID3 analysis [45] was initiated with mintotal =3000, minexp =5, minnumber=5, and internal batch effect using the plates ID as input. For the outlier identification we used probthr =0.001 and outlog=2. Then, the transcription distance was calculated using the compdist() function, followed by clustexp() using the following parameters: clustnr= 30, bootnr =50, samp=1000, metric = pearsonto, sat= TRUE, FUNcluster = “hclust” to identified the initial number of clusters present in our dataset. We used the plotjaccard() function to identify the saturation point and re-calculated the clustexp() function using cln=8 and sat=FALSE. After RaceID3 benchmarking, we performed VarID analysis [46] to infer the local neighborhoods of cells in the same. To determine the k-nearest neighbors for each cell in gene expression space, and test if the links are supported by a negative binomial joint distribution of gene expression we applied the pruneKnn function with the following parameters: d= gene expression values from RaceID3 dataset, pcaComp=100, regNB=TRUE, batch=plate ID name, knn=10, alpha =1, ngenes=2000 and links with probabilities lower than pvalue < 0.01 were removed. To visualize the VarID analysis, we performed Louvain clustering and UMAPs reduction for cluster visualization were generated using compumap() function. To increase our HSPC cluster resolution we removed the erythrocytes and myeloid cells clusters from our single-cell object using the reduceset() function. Differentially expressed genes were identified by the differential gene expression function diffexpnb() from the RaceID3 (v0.1.4) algorithm with multiple testing corrected by the Benjamini-Hochberg method set significance as p<0.01. ClurterProfiler software version 4.0 (https://doi.org/10.1016/j.xinn.2021.100141) was run to evaluate the biological pathways associated with each of differentially expressed genes (DEG) from the VarID identified clusters. The parameters applied were: all identified transcript set as universe, the DEG form each target cluster set as gene target list and enrichment strategy set either as enrichGO() or enrichKEGG(). Significance was scored by adjusted Benjamin-Hochberg with p-value = 0.01 and qvalue = 0.05.

### Whole mount in situ hybridization (WISH)

*In situ* hybridization was performed using a standard protocol [47]. *In situ* probes were generated by PCR amplification using a cDNA or plasmid template followed by reverse transcription with digoxigenin-linked nucleotides. Primer sequences for all WISH probes used in this paper are provided in Supplementary Table 1.

### Transgenesis

Transgenic lines were established as previously described [48]. The *mpeg1:TagBFP* line was generated by PCR amplification of the pCAG-dGBP1-TagBFP plasmid, D-TOPO cloning into pME Gateway, and LR Gateway Reaction to generate constructs with the *mpeg1* promoter driving *TagBFP,* followed by an SV40 polyA signal sequence, all flanked by Tol2 integration sites. The pCAG-dGBP1-TagBFP plasmid was a gift from Connie Cepko [49] (Addgene plasmid #80086). Calreticulin overexpression constructs were generated by PCR amplification off of genomic DNA, followed by D-TOPO cloning and LR gateway reactions as noted before. The fidelity of all constructs was confirmed by sequencing prior to injection.

### Microscopy and image analysis

Time-lapse microscopy was performed using a Yokogawa CSU-X1 spinning disk mounted on an inverted Nikon Eclipse Ti microscope equipped with dual Andor iXon EMCCD cameras and a climate controlled (maintained at 28.5°C) motorized x-y stage to facilitate tiling and imaging of multiple specimens simultaneously. Screening of injected constructs and imaging of WISH embryos was performed using a Nikon SMZ18 stereomicroscope equipped with a Nikon DS-Ri2 camera. All images were acquired using NIS-Elements (Nikon), blinded, and processed using Imaris (Bitplane). Specimens were mounted in 0.8% LMP agarose with tricaine (0.16 mg/ml) in glass bottom 6-well plates and covered with E3 media containing tricaine (0.16 mg/ml).

### Flow cytometry

To collect kidney marrows, adult zebrafish (2 to 9-months-old) were anaesthetized with 0.02% tricaine in fish water and dissected under a Leica MZ75 light microscope. The soft tissue of the kidney marrow was placed in cold 0.9x DPBS (Gibco) with 2% fetal bovine serum (FBS, Gemini Bio-Products) and 1 USP units/mL heparin (Sigma), and then mechanically dissociated by repeated pipetting into single cell suspension, and passed through a 40-μm nylon mesh 5-10 minutes prior to analysis. To collect embryonic samples, embryos were chopped with a razor blade in cold PBS and then incubated in Liberase (Roche) for 20 minutes at 37°C before filtering the dissociated cells through a 40 μm filter and transferring to 2% FBS. For samples stained for Calreticulin, Liberase was not added. Cells were blocked in 5% Normal Goat Serum + 0.5% Sodium Azide in cold PBS for 20 minutes, washed, and incubated with a Chicken anti-Calreticulin antibody (abcam ab94935, 1:1000) for 30 minutes on ice. Samples were then washed and incubated with a Goat anti-Chicken Alexa Fluor 488 antibody (Invitrogen A11039, 1:1000) for 25 minutes at room temperature, protected from light. CellROX Deep Red (Invitrogen C10422) and AnnexinV-FITC (BD Biosciences) staining was performed according to manufacturer instructions. We used 3nM DRAQ-7 for live/dead stain (Abcam). Flow cytometric analysis was performed on a BD FACSAria II with special order 445nm laser for CFP detection when using Zebrabow (BD Biosciences). Gates were drawn using negative and isotype matching controls. Data were analyzed with FlowJo software version 10.

### Zebrabow color labelling

At 24 hours post fertilization (hpf) embryos were transferred to 6-well plates at a density of 25-35 embryos per well and treated with 15 μM 4-hydroxytamoxifen (4-OHT) for 3-5 hours in the dark at 28.5°C.

### Zebrabow analysis

Color barcodes from Zebrabow kidney marrow samples were quantified using previously published pipelines [7] adapted to a Python-based interface. The granulocytic color output was chosen as a read out of clonal changes as these cells have a short half-life and reflect changes in the HSPC clonal output in a timely manner. Only zebrafish with greater than 75% recombination efficiency were processed. All samples were blinded prior to analysis.

### Drug treatment

Drugs were added to embryo E3 media. MEK1/2 inhibition was done using PD98059 (Sigma) at 48 hpf at a concentration of 15 μM. Diphenylene iodonium (Sigma) was added to embryos at 48 hpf at a concentration of 100 μM.

### Morpholino injections

Morpholinos (GeneTools) were resuspended to 300 μM in nuclease free water, heated to 65°C for 5 minutes, and kept at room temperature. Embryos were injected with 1 nanoliter into the yolk at the 1-4 cell stage with either 3 ng (*calr, calr3a, calr3b, std*) or 5 ng (*irf8, std*) of morpholino. Morpholino sequences are listed in Supplemental Table 2.

### Liposome injection

Zebrafish embryos were dechorionated and anesthetized with tricaine (0.16mg/ml) on flat agarose disks. Approximately 1.5 nanoliters of liposomes loaded with either clodronate or PBS (Liposoma) were injected directly into circulation into either the caudal vein (28 hpf) or the duct of Cuvier (48 hpf, 72 hpf).

### Few-cell proteomics

Cell pellets were submitted after FACS sort for mass spec analysis. Cell pellets were resolubilized in pure water and undergoing mPOP procedure [50], briefly, frozen cells in water straight from −80C freezer were put into heated 96 well plate shaker at 95 C for 5 min. After cells were chilled to room temperature trypsin was added for 3 hours digest procedure at 38 C. Digested proteins were labeled with TMT11plex (Tandem Mass Tags) labels (Thermo-Fisher, Germany) according to the previously published procedure SCOPE-MS [18]. After labeling procedure all samples were quenched with 5% hydroxyl solution and pulled together as one sample for further mass spectrometry analysis. Two consecutive LC-MS/MS experiments were performed on HFX Orbitrap (Thermo-Fisher, CA) equipped with UltiMate 3000 HPLC Nano tandem pump (Thermo-Fisher, CA). Peptides were separated onto PharmaFluidics (Belgium) trapping column followed by elution into 50 cm PharmaFluidics analytical column. Separation was achieved through applying a gradient from 5–27% ACN in 0.1% formic acid over 180 min at 200 nl/min. Electrospray ionization was enabled through applying a voltage of 2 kV using a home-made electrode junction at the end of the microcapillary column and sprayed from stainless steel 4 cm needles (PepSep, Denmark). The HFX Orbitrap instrument was operated in data-dependent mode for the mass spectrometry methods. The mass spectrometry survey scan was performed in the Orbitrap in the range of 400 –1,400 m/z at a resolution of 1.2 × 105, followed by the selection of the twenty most intense ions (TOP10) for HCD-MS2 fragmentation in the Orbitrap using a precursor isolation width window of 0.7 Th, AGC setting of 50,000, and a maximum ion accumulation of 200 ms. Singly charged ion species were not subjected to HCD fragmentation. Normalized collision energy was set to 34 V and an activation time of 1 ms. Ions in a 10 ppm m/z window around ions selected for MS2 were excluded from further selection for fragmentation for 90 seconds. Raw data were submitted for analysis in Proteome Discoverer 2.4. (Thermo-Fisher, CA) software. Assignment of MS/MS spectra were performed using the Sequest HT and Byonic v3.5 (Protein Metrics, CA) algorithms by searching the data against a protein sequence database including all entries from Zebrafish database and other known contaminants such as human keratins and common lab contaminants. Sequest HT and Byonic searches were performed using a 10 ppm precursor ion tolerance and requiring each peptides N-/C termini to adhere with Trypsin protease specificity, while allowing up to two missed cleavages. 6-plex TMT tags on peptide N termini and lysine residues (+229.162932 Da) was set as static modifications while methionine oxidation (+15.99492 Da) was set as variable modification and deamidation of Asparagine and Glutamine amino acids. An MS2 spectra assignment false discovery rate (FDR) of 1% on protein level was achieved by applying the target-decoy database search. Filtering was performed using a Percolator [51] (64bit version).

For quantification, a 0.02 m/z window centered on the theoretical m/z value of each the six reporter ions and the intensity of the signal closest to the theoretical m/z value was recorded. Reporter ion intensities were exported in result file of Proteome Discoverer 2.4 search engine as an excel tables. The total signal intensity across all peptides quantified was summed for each TMT channel, and all intensity values were adjusted to account for potentially uneven TMT labeling and/or sample handling variance for each labeled channel. All further data evaluation for statistically differential analysis were performed on in-house made R package based on Bioconductor programs (https://www.bioconductor.org/).

### Immunofluorescence Staining

Mouse embryos were harvested at 14.5 days post conception. Fetal livers were dissected and stored in 4% paraformaldehyde at 4°C overnight. Samples were dehydrated in 30% sucrose for 8 hours and embedded in OCT. Cryosections were collected at a thickness of 10 microns. Slides were washed three times in PBS + 0.1% Triton, blocked with 5% normal goat serum and 1% bovine serum albumin for two hours, and stained overnight with rabbit anti-c-Kit (Cell Signaling Technology #3074, 1:400) and rat anti-F4/80 (abcam ab6640, 1:100). Slides were washed five times with PBS + 0.1% Triton and stained for two hours with Goat anti-rabbit Alexa Fluor 488 (Invitrogen A11008, 1:500), Goat anti-rat Alexa Fluor 568 (Invitrogen A11077, 1:500) and DAPI (abcam ab228549, 1:1000). Slides were washed 5 times with PBS + 0.1% Triton and coverslips were mounted with ProLong^TM^ Diamond Antifade Mountant.

### Zebrafish EdU Labeling

Embryonic circulation was injected at 72 hpf with 1 nanoliter of 500μM EdU. Embryos were kept at 4°C for 1 hour, fixed in 4% paraformaldehyde for 1 hour, permeabilized with 0.1% Triton for 20 minutes at room temperature and labeled with Alexa Fluor 647 using the Click-iT reaction (Thermo Fisher) for 30 minutes according to manufacturer instructions. Embryos were then washed with PBS+0.5% Triton and blocked for one hour with 10% Normal Goat Serum, 0.5% Bovine Serum Albumin, 0.5% Triton. Samples were incubated with Rat anti-mCherry Alexa Fluor 594 (Invitrogen M11240, 1:200) for one hour at room temperature and washed 5 times with PBS+0.5% Triton.

### Parabiosis

Parabiotic zebrafish generated as previously described [52]. Briefly, zebrafish embryos were injected with morpholinos, and incubated until the 1,000-cell stage. Pairs were fused by embedding in 4% methylcellulose, immersion in high-calcium Ringer’s solution containing penicillin and streptomycin, and surgical wounding and connection of partner embryos at the animal pole.

### Acridine Orange Staining

Acridine Orange dye (Invitrogen A3568) was used at a final concentration of 3μg/mL in E3 embryo media. Embryos were incubated with dye for 30 minutes, washed twice with E3, and mounted for imaging.

### Statistical analysis

Graphs and statistical analyses were done with Prism (Graphpad), Excel (Microsoft), and RStudio. For all graphs, error bars indicate mean +/− standard deviation. P values were obtained with two-tailed Student’s t-¬test or One-way ANOVAs for all analyses as indicated.

## Supplementary Materials

Figs. S1 to S5

Tables S1 to S2

Movies S1 to S3

## Acknowledgments

We gratefully thank the Boston Children’s Hospital veterinary staff, the BCH flow cytometry core, Single Cell Discoveries, and the Harvard Center for Mass Spectrometry. We also thank A. Han for her assistance with fetal mouse dissection, B. Miller for her help preparing cryosections, and colleagues for critical reading of the manuscript. S. Wattrus thanks P. Chen for her continued support in all things.

## Funding

National Institutes of Health grants 1F31HL149154-01 (SJW); 5T32HL007574-40 (CPR); K01DK111790 (EJH); P01HL131477, P01HL032262, U54DK110805, R24DK092760, R24OD017870, U01HL134812, and R01HL144780-01 (LIZ). The Edward P. Evans Foundation (LIZ). Alex’s Lemonade Stand Fund (LIZ). LIZ is a Howard Hughes Medical Institute Investigator.

## Author contributions

SJW designed the study, funded the project, performed experiments, managed the project, interpreted the data, and wrote the manuscript. MLS. Performed experiments, managed the project, interpreted the data, and edited the manuscript. CPR, EJH, and BB performed experiments and interpreted the data. LIZ designed the study, funded and supervised the project, interpreted the data, and edited the manuscript. All authors reviewed the manuscript.

## Competing interests

LIZ is a founder and stockholder of Fate Therapeutics, CAMP4Therapeutics, and Scholar Rock. All other authors declare that they have no competing interests.

## Data and materials availability

All data are available in the main text or the supplementary materials. The scRNA-seq data have been submitted to the NCBI Gene Expression Omnibus and accession numbers will be forwarded upon receipt. Proteomic data has been submitted to the MassIVE database and PRIDE repository and accession numbers will be available upon receipt.

