## Supplemental Information for "Quality assurance of hematopoietic stem cells by macrophages determines stem cell clonality"

Contains 5 Supplemental Data Figures, 3 Supplemental Videos, and 2 Supplemental Tables

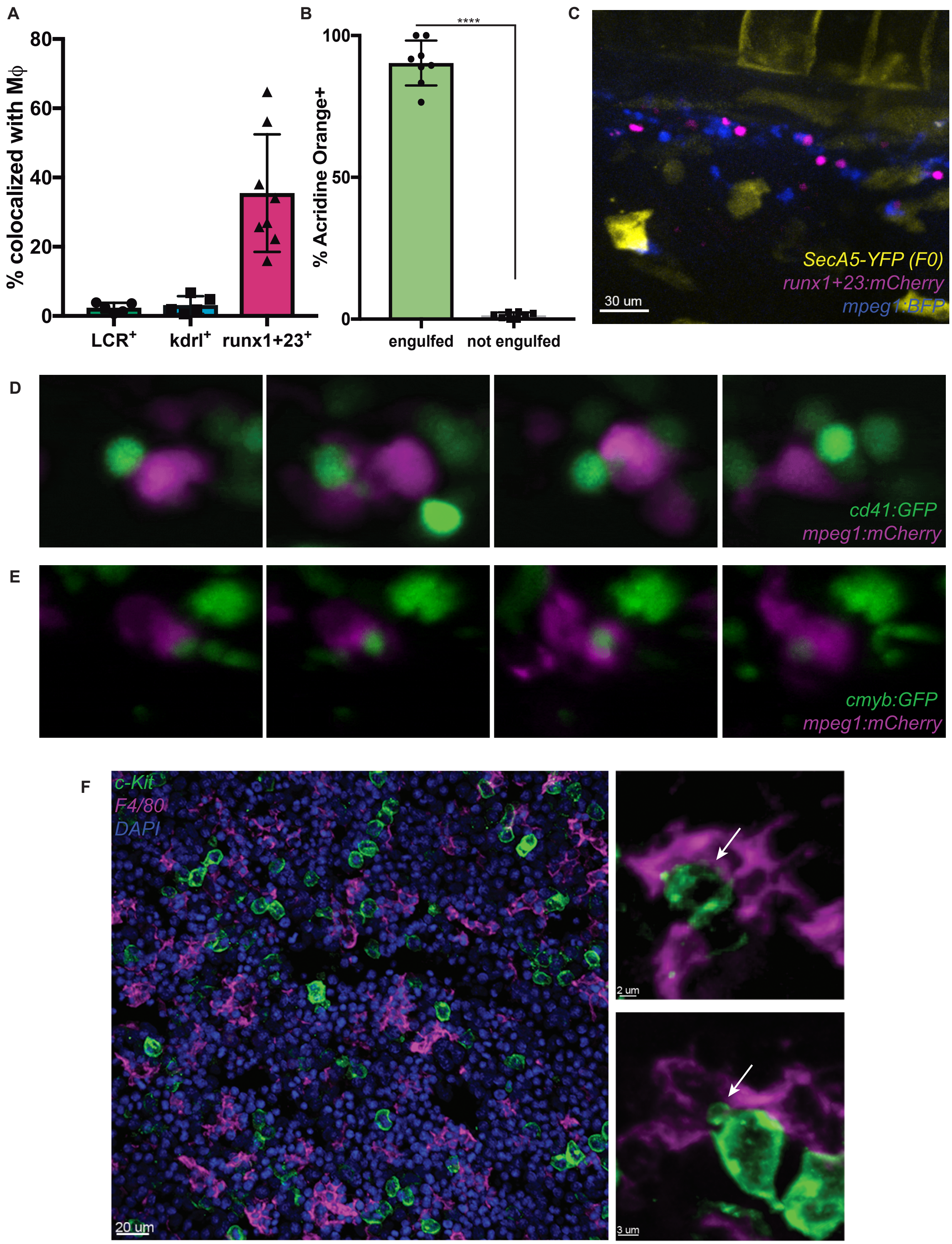


**Supplemental Figure 1**

**Fig. S1. Macrophages specifically interact with HSPCs.**

(**A**) Quantification of macrophage contacts with *kdrl*$^{+}$ endothelial cells and *LCR*$^{+}$ erythroid cells in the CHT shows minimal levels of interaction compared to the fraction of *runx1+23*$^{+}$ HSPCs engaged by macrophages. Mean +/- s.d. (**B**)(**C**) The only Acridine Orange$^{+}$ or AnnexinV-YFP$^{+}$ apoptotic HSPCs in the CHT are those which have been engulfed by macrophages. At baseline, there are no apoptotic HSPCs. Mean +/- s.d., Unpaired t test; ****$P<0.0001$. (**D**)(**E**) Cells labeled by *cd41:GFP* and c*myb:GFP* - transgenes which also mark HSPCs – also undergo intimate interactions with macrophages in the CHT which involve fragments of cytoplasmic fluorescent material exchanged or full cell engulfment. (**F**) Immunofluorescence staining of murine fetal liver at E14.5 reveals that approximately 33% cKit$^{+}$ hematopoietic progenitors are in contact with F4/80$^{+}$ macrophages, including interactions in which pieces of or entire cKit$^{+}$ cells appear to be taken by macrophages.

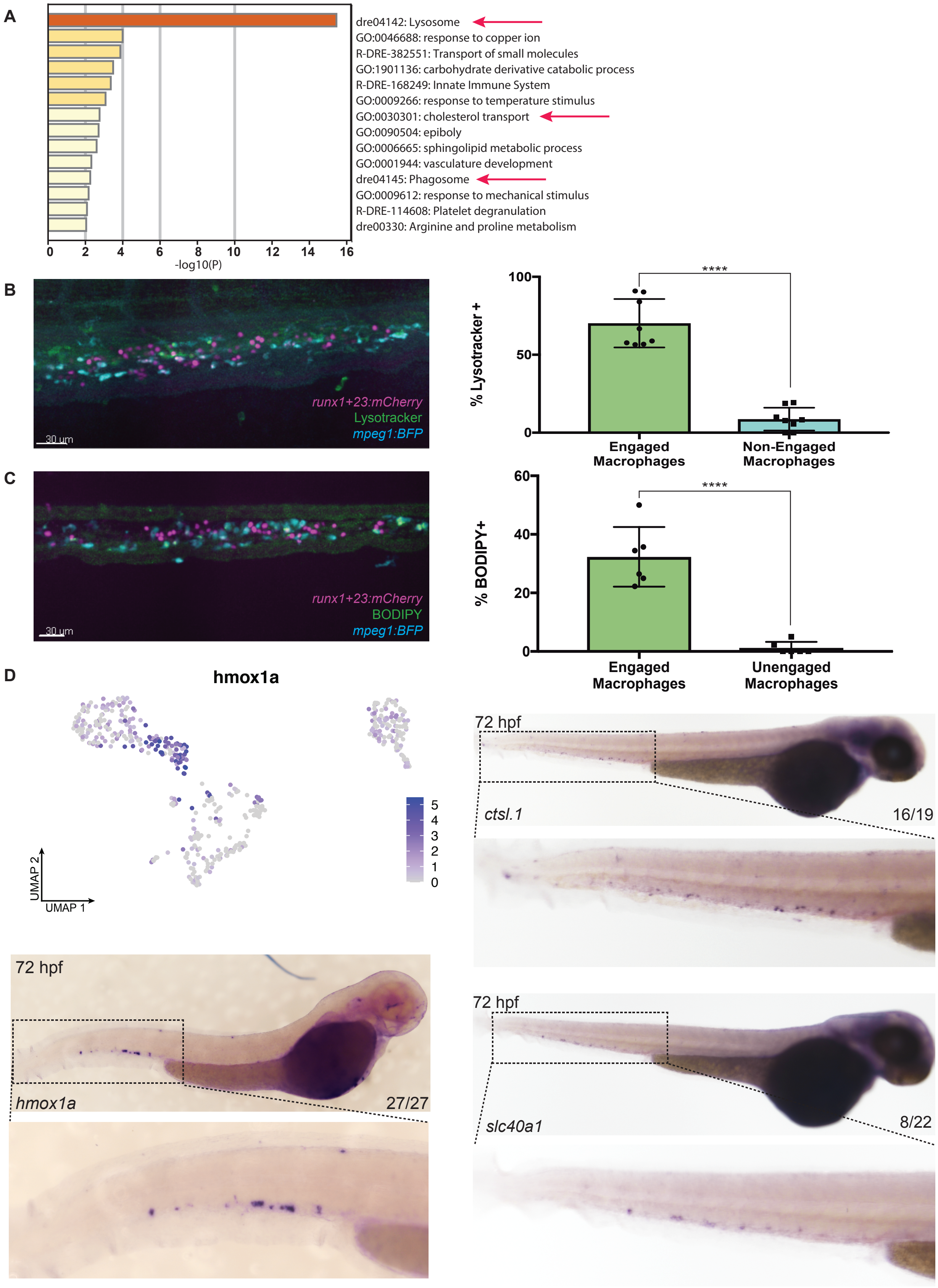


**Supplemental Figure 2**

**Fig. S2. Interacting macrophages are a distinct population.**

(**A**) GO-term enrichment for genes highly enriched in the macrophages which have taken up fragments of or entire HSPCs. Arrows indicate GO-terms of note: lysosome, cholesterol transport, and phagosome. (**B**) Lysotracker dye marks macrophages which interact with HSPCs in the CHT. Scale bar indicates 30μm. Mean +/- s.d., Unpaired t test; ****$P<0.0001$. (**C**) The cholesterol mimic, BODIPY, is specifically taken up by macrophages which interact with HSPCs. Scale bar indicates 30μm. Mean +/- s.d., Unpaired t test, ****$P<0.0001$. (**D**) UMAP of single-cell mRNA-seq for 72 hpf niche macrophages shows enrichment for the marker *hmox1a* in the subset of macrophages which interact with HSPCs. Whole mount *in situ* hybridization confirms expression of *hmox1a, ctsl.1,* and *slc40a1* in a subset of macrophages in the CHT at 72 hpf. Spectral scale indicates z score.

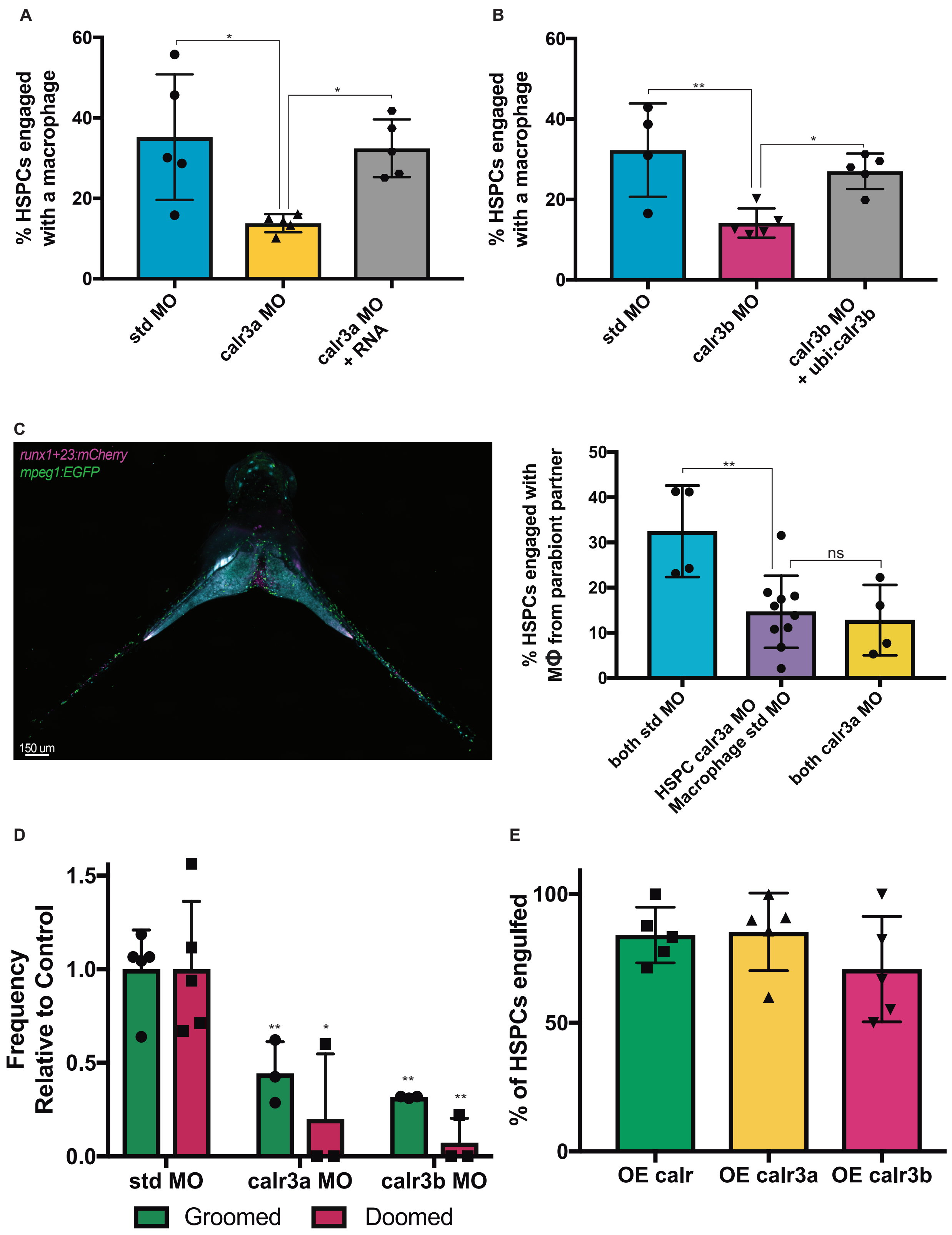


**Supplemental Figure 3**

**Fig. S3. Calreticulin knock-down can be rescued and affects grooming and dooming.**

(**A**)(**B**) The fraction of HSPCs interacting with macrophages is reduced by injection of either the *calr3a* or *calr3b* morpholino but can be rescued by co-injection of either mRNA or DNA constructs encoding ubiquitous Calreticulin expression. Mean +/- s.d., One-way ANOVA; *P<0.05, **P<0.01. (**C**) Example and quantification of parabiotic fusion of *runx1+23:mCherry* and *mpeg1:EGFP* embryos. HSPCs with *calr3a* knock-down have reduced interactions with unperturbed macrophages. Mean +/- s.d., One-way ANOVA with Dunnett's multiple comparisons test; **P<0.01. (**D**) Knock-down of *calr3a* or *calr3b* reduces the fraction of HSPCs that are groomed or doomed by macrophages. Mean +/- s.d., One-way ANOVA; *P<0.05, **P<0.01. (**E**) Most HSPCs overexpressing surface translocated forms of *calr* (84%), *calr3a* (85%), or *calr3b* (71%) are fully engulfed by macrophages.

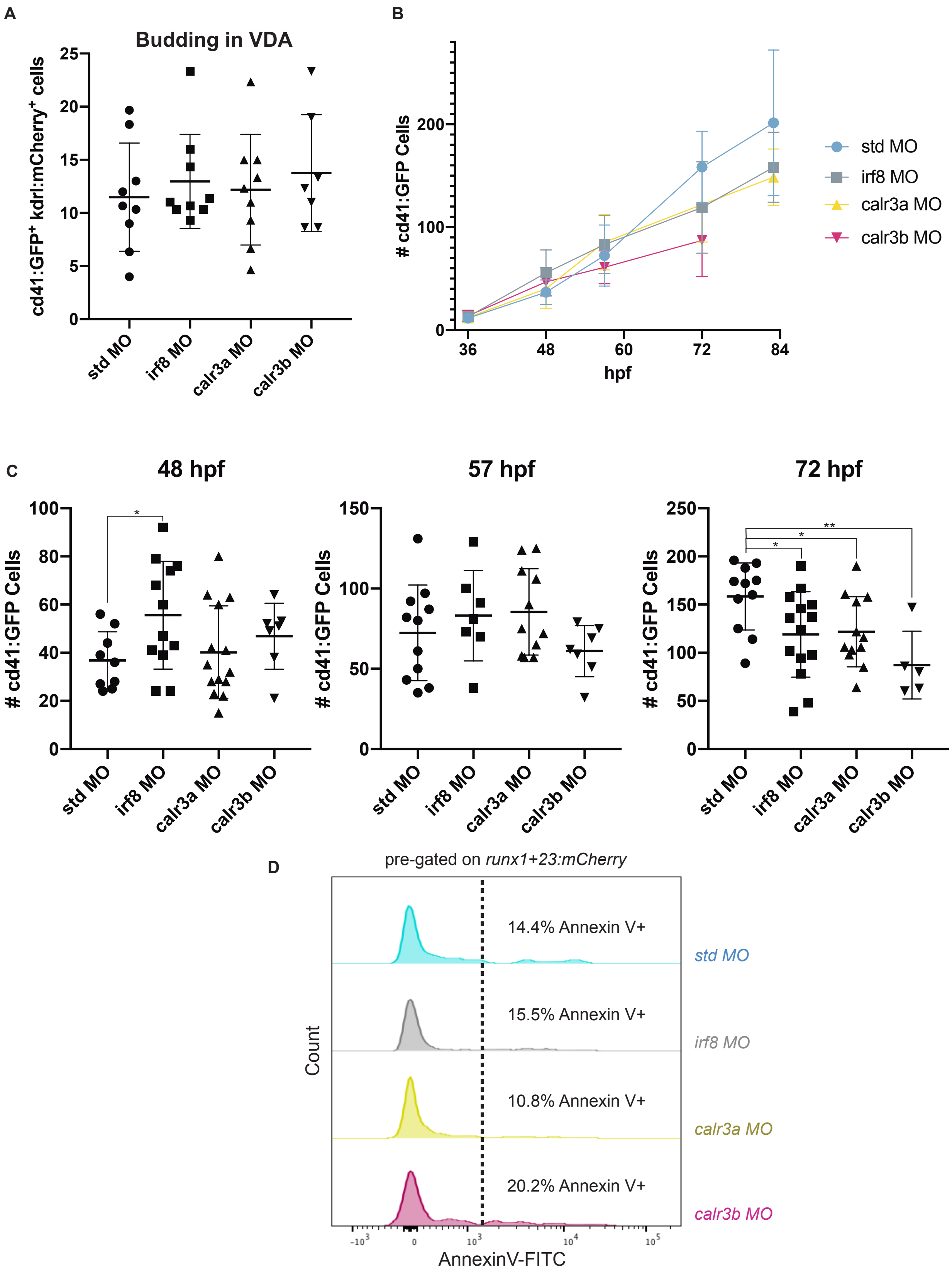


**Supplemental Figure 4**

**Fig. S4. Calreticulin knock-down reduces the number of HSPCs in the CHT.**

(**A**) Quantification of *kdrl:mCherry;cd41:GFP*$^+$ cells in the floor of the dorsal aorta from 32-46 hpf reveals no significant change in the number of emerging HSPCs in embryos injected with the calr3a, calr3b, or irf8 morpholino. Mean +/- s.d. (**B**) Serial imaging of *cd41:GFP* embryos identify reduced numbers of HSPCs in the CHT at 3 dpf in the absence of macrophage interactions. Similar numbers of cells emerge from the aorta and initially seed the CHT, but embryos injected with the *irf8*, *calr3a*, or *calr3b* morpholinos exhibit a deficit from 60 – 84 hpf. Mean +/- s.d. (**C**) Quantitation of *cd41:GFP*$^+$ cells in the CHT at 48 hpf, 57 hpf, and 72 hpf. Mean +/- s.d., Unpaired t test; *P<0.05, **P<0.01. (**D**) Annexin V staining and flow cytometry after injection with either the *calr3a, calr3b,* or *irf8* morpholino finds similar fractions of *runx1+23*$^+$ cells undergoing cell death compared to control morphants.

A

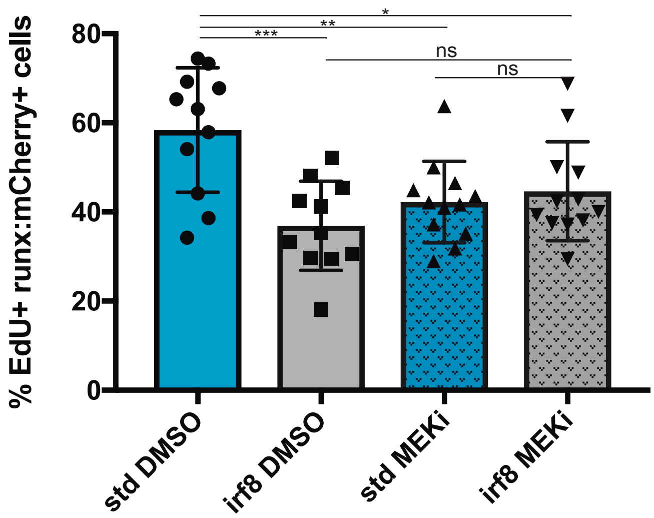


B

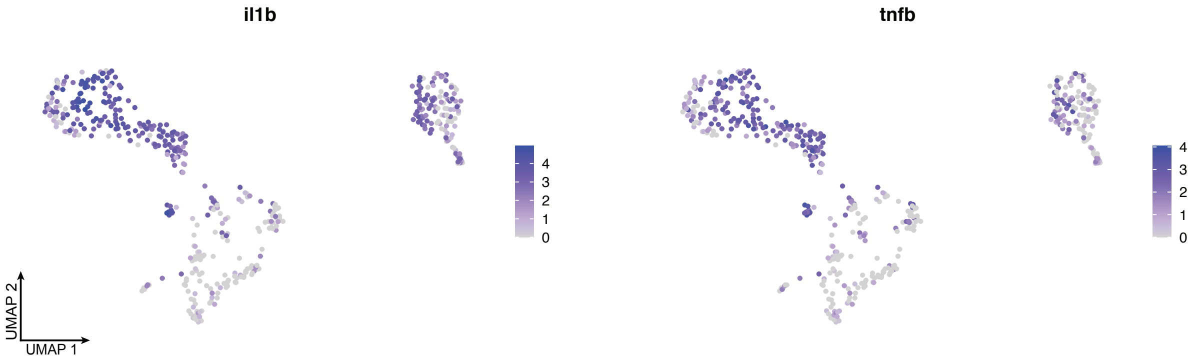


Supplemental Figure 5

**Fig. S5. Macrophages express pro-proliferative signals capable of ERK/MAPK signaling**

(**A**) Both macrophage depletion and MAPK inhibition reduce HSPC proliferation. The combination of macrophage depletion and MAPK inhibition has no additional impact on HSPC divisions. Mean +/- s.d., Unpaired t test; *$P<0.05$, **$P<0.01$, ***$P<0.001$. (**B**) Macrophages in the CHT express *il1b* and *tnfb*, signals that have previously been noted for their capacity to induce HSPC division. Spectral scales report z score.

### Table S1. Primer sequences used in this study

| | |
|---|---|
| *hmox1a* ISH fwd | 5' CATTAACCCTCACTAAAGGGAACAGGACTTGGAGCACTTCTTCGG 3' |
| *hmox1a* ISH rev | 5' TAATACGACTCACTATAGGGAGACAACTCAAAGCGTTTATGGACAG 3' |
| *ctsl.1* ISH fwd | 5' CATTAACCCTCACTAAAGGGAAGATCAGAAACAGTGTGGATCATGC 3' |
| *ctsl.1* ISH rev | 5' TAATACGACTCACTATAGGGCCTAGACAGGGGCATTAAAAATGAAAACAG 3' |
| *slc40a1* ISH fwd | 5' CATTAACCCTCACTAAAGGGAAAAAACCTGTCGCCGAAGTTCAC 3' |
| *slc40a1* ISH rev | 5' TAATACGACTCACTATAGGGATAATCCTCCCACTGCGATGG 3' |
| *calr* fwd | 5' CACCATGACTGCGTTATCCCTACTGTTTATG 3' |
| *mTagBFP* fwd | 5' CACCATGAGCGAGCTGATTAAGGAGAACA 3' |
| *mTagBFP* rev | 5' TTAATTAAGCTTGTGCCCCAGTTTGCT 3' |
| *calr*NOKDEL NO STOP rev | 5' GAGTTTAGAGTCTGTTTCCTCCTCCTC 3' |
| *calr3a* fwd | 5' CACCATGCGGATCACTGCTGCA 3' |
| *calr3a* rescue fwd | 5' CACCATGAGAATAACAGCCGCCGTGTGCTTTATTTCTGCACTGGC 3' |
| *calr3a* rev | 5' GATCATCTAGACTACAATTCATCTTTAGGGAGCGCATCAT 3' |
| *calr3a*NOKDEL NO STOP rev | 5' AGGGAGCGCATCATCCTCA 3' |
| *calr3b* fwd | 5' CACCATGCAAATTTCATTATTACAGTTAATTTCGGCT 3' |
| *calr3b* rev | 5' GATCATCTAGACTACAGTTCGTCTTTCTGAAGCACATC 3' |
| *calr3b*NOKDEL NO STOP rev | 5' CTGAAGCACATCCTCGTCTCC 3' |

**Table S2. Sequences for morpholinos used in this study**

| | |
|---|---|
| standard control | 5’ CCTCTTACCTCAGTTACAATTTATA 3’ |
| *irf8* | 5’ TCAGTCTGCGACCGCCCGAGTTCAT 3’ |
| *calr* | 5’ AACAGTAGGGATAACGCAGTCATCT 3’ |
| *calr3a* | 5’ GCAGCAGTGATCCGCATCTCTGCAC 3’ |
| *calr3b* | 5’ AAAGAAATGACTCCCGCACGCTCGC 3’ |

**Movie S1. Macrophages take up fluorescent material from HSPCs.**

Video of Fig. 1A; an *mpeg1:mCherry*$^{+}$ crawls over the surface of a *runx1+23:EGFP*$^{+}$ HSPC at 72 hpf and sucks up cytoplasmic EGFP material.

**Movie S2. Macrophages may groom or doom HSPCs in the CHT niche.**

Video showing *mpeg1:EGFP* macrophages interacting with *runx1+23:mCherry* HSPCs in the CHT at 72 hpf. Arrows indicate fully engulfed "doomed" HSPC that is degraded, and an HSPC that is "groomed" and has fragment of material taken up. Circle indicates HSPC about to divide.

**Movie S3. HSPCs only undergo apoptosis upon dooming.**

Video showing a *runx1+23:mCherry*$^{+}$ HSPC being doomed by an *mpeg1:BFP*$^{+}$ macrophage in the CHT at 72 hpf. The cell does not become Acridine Orange$^{+}$ until it is fully engulfed.
